# Measuring thousands of single vesicle leakage events reveals the mode of action of antimicrobial peptides

**DOI:** 10.1101/2021.08.13.455434

**Authors:** Kareem Al Nahas, Marcus Fletcher, Katharine Hammond, Christian Nehls, Jehangir Cama, Maxim G Ryadnov, Ulrich F Keyser

## Abstract

Host defense or antimicrobial peptides hold promise for providing new pipelines of effective antimicrobial agents. Their activity quantified against model phospholipid membranes is fundamental to a detailed understanding of their structure-activity relationships. However, existing characterization assays lack the resolution necessary to achieve this insight. Leveraging a highly parallelized microfluidic platform for trapping and studying thousands of giant unilamellar vesicles, we conducted quantitative long-term microscopy studies to monitor the membrane-disruptive activity of archetypal antimicrobial peptides with a high spatiotemporal resolution. We described the modes of action of these peptides via measurements of the disruption of the vesicle population under the conditions of continuous peptide dosing using a range of concentrations, and related the observed modes with the molecular activity mechanisms of these peptides. The study offers an effective approach for characterizing membrane-targeting antimicrobial agents in a standardized manner, and for assigning specific modes of action to the corresponding antimicrobial mechanisms.

## Introduction

The COVID-19 pandemic has refocused the world’s attention on the dramatic consequences of failing to contain infectious diseases. However, the “silent” pandemic of antimicrobial resistance (AMR) continues unabated and remains mostly out of the public eye. The spread of multidrug resistant bacteria, combined with both economic^1^ and scientific^2^ challenges in drug development, has led to a situation where common infections are becoming untreatable. There is a dire need to provide new pipelines of antimicrobial agents exhibiting mechanisms which may be less subject to acquired resistance.

Molecular agents disrupting microbial membranes are often advantageous over traditional antibiotics targeting individual intracellular processes in microbial cells.^3^ Their activity is not necessarily confined to a single target, does not require reaching the cytoplasm, but is not limited to membrane disruption either, allowing them to tackle a microbial cell as a whole. Such agents are of particular interest for treating Gram-negative bacteria whose double membranes continue being a formidable barrier for antibiotics with intracellular targets. ^4,5^ Host defence or antimicrobial peptides (AMPs) are typical representatives of membrane-active antimicrobials. These peptides favour attack on bacterial cell surfaces demonstrating high efficacy against multidrug- and pandrug-resistant bacteria *in vitro*.^6^ Yet, the translation of these agents and treatments into clinical use has not been efficient, suggesting a discordance between *in vitro* tests and clinical trials when attempting to balance out the efficacy, stability and toxicity of these peptides. As the field has also recently noted,^7,8^ better and standardized characterization platforms are needed to facilitate a deeper and more systematic understanding of the antimicrobial mechanisms of these peptides. This will support more predictive structure-activity outcomes and hence facilitate the efficient clinical translation of AMPs.

When developing these tools, the terms “mechanism” and “mode of action” are often used interchangeably. In this study we distinguish between the two in that “mechanism” refers to the dynamics governing interactions between peptides and phospholipids at the molecular level. This mechanistic understanding can be elucidated by employing techniques such as Molecular Dynamics (MD) simulations,^10^ oriented circular dichroism spectroscopy^11^ and atomic force microscopy (AFM).^12^ By “mode of action” we mean the activity of the peptides at the cell population level using either live cells or model membranes such as unilamellar vesicles. This may involve the kinetics of membrane disruption as determined by the distribution of leakage timepoints within a population of vesicles. With a molecular mechanism underpinning a mode of action, we chose to make this distinction explicitly.

Peptide-membrane interactions are postulated to occur via the so-called “two state model”; the surface state, during which peptides bind to membrane surfaces, and the insertion state when bound peptides insert into the membrane forming pores and channels.^13^ In this model the point of transition between the two states is defined by reaching a threshold concentration of bound peptide after a certain lag period. The insertion state can be followed by pore expansion causing critical membrane disruption and eventually leading to cell lysis.^12,14^

Leakage studies using reconstituted synthetic unilamellar vesicles provide a straightforward probe of the antimicrobial activity of membrane active AMPs. Typically, a fluorescent probe is encapsulated in the lumen of a phospholipid vesicle, and a stable (non-decaying) fluorescence signal from the vesicle indicates that the lipid membrane remains intact. Conversely, a signal decay (after controlling for photobleaching) suggests that the encapsulated fluorophore leaks out due to a compromised membrane.^15^ Monitoring peptide-induced leakage kinetics can reveal the “mode of action” of the peptide, which in turn can help interpret a particular permeabilization “mechanism”.^16,17^ In this vein, most previous studies used a suspension of large unilamellar vesicles (LUVs; diameters spanning 100–1000 nm) for monitoring peptide activity. In such cases, the user measures the average fluorescence values of the LUVs as a single population, and hence such experiments provide information on the average leakage kinetics of the vesicle population as a whole. Consequently, the coarse-grained resolution of the extracted information only enables minimal interpretation of the mode of action - it does not allow the investigation of heterogeneity in the mode of action at the single vesicle level. To enhance the quality and level of detail extracted from these experiments, Giant unilamellar vesicles (GUVs), which can be studied individually, are used instead.^18^ In such studies, a handful of vesicles (diameters >1000 nm) can be directly visualized using fluorescence microscopy enabling the monitoring of the peptide-induced leakage kinetics of individual GUVs with a defined spatiotemporal resolution. However, such methods require laborious manual handling, and suffer from a lack of control over experimental conditions and low throughput. To overcome these challenges, we previously devised a bespoke microfluidic platform that integrated high-throughput GUV formation and immobilization on-chip in parallel trapping arrays.^19^ This format then enabled the continuous administration of different peptides at a set dose in a parallelized manner on thousands of trapped vesicles. ^20^ This microfluidic platform is thus perfectly suited to investigating the mode of action of membrane-active agents on a population of thousands of model membranes, while maintaining single-GUV and high spatiotemporal resolution. In this work, we leverage our ability to monitor the response of thousands of individual GUVs to a range of AMPs. We optimize a bespoke microfluidic platform into an effective high-throughput device, dubbed a GUV studio, and demonstrate its applicability to elucidate antimicrobial modes of action. We relate the modes of action to the timing of dye leakage from the GUVs after applying a defined peptide dosage and with molecular mechanisms elucidated by atomic force microscopy. By identifying and quantifying modes of disruption or leakage and correlating them with molecular mechanisms, we provide information that can inform appropriate dosage and administration regimens for antimicrobial agents.

## Results and Discussion

We employed an optofluidic leakage assay to monitor single vesicle leakage events (LEs) utilizing the GUV studio depicted in figure 6. The platform allows the user to continuously dose several populations of GUVs (each containing a membrane impermeable dye) with an antimicrobial agent at different concentrations in parallel over 7.5 hours. A vesicle LE is defined as the >50% loss of fluorescent dye from the vesicle lumen. Importantly, within each experiment, all the trapped populations of GUVs were assembled on the same device. The source of all the GUVs in an experiment was hence identical.

To allow direct comparisons, the experimental parameters and conditions were identical for all the peptides studied, with peptide sequence being the only variable. To demonstrate the approach, two series of representative antimicrobial peptides were selected. One series consisted of bacteriocin-derived peptides including a four-helix bacteriocin epidermicin (NI01), its all-D analogue (D-NI01), an arginine mutant (R-NI01) and three 2-helix substructure derivatives (Figure 1).^9^ Another series was comprised of well-studied host defence peptides discovered in different organisms, including melittin (a bee venom peptide),^21^ an amphibian peptide magainin 2 (from *Xenopus laevis*),^22^ an antibiotic alamethicin from the fungus *Trichoderma viride*,^23^ a last-resort antibiotic polymyxin B (from *Bacillus polymyxa*),^24^ and a human β-defensin (hβD-3-l). ^25,26^ The response of a GUV population to defined concentrations of each peptide was monitored with single GUV resolution. Four different peptide concentrations were run in parallel to probe concentration dependent effects. The mode of action was determined by measuring the time distribution of single GUV LEs and the percentage survival of a GUV population at the end of an experiment as summarized in the corresponding heat maps.

**Figure 1:**
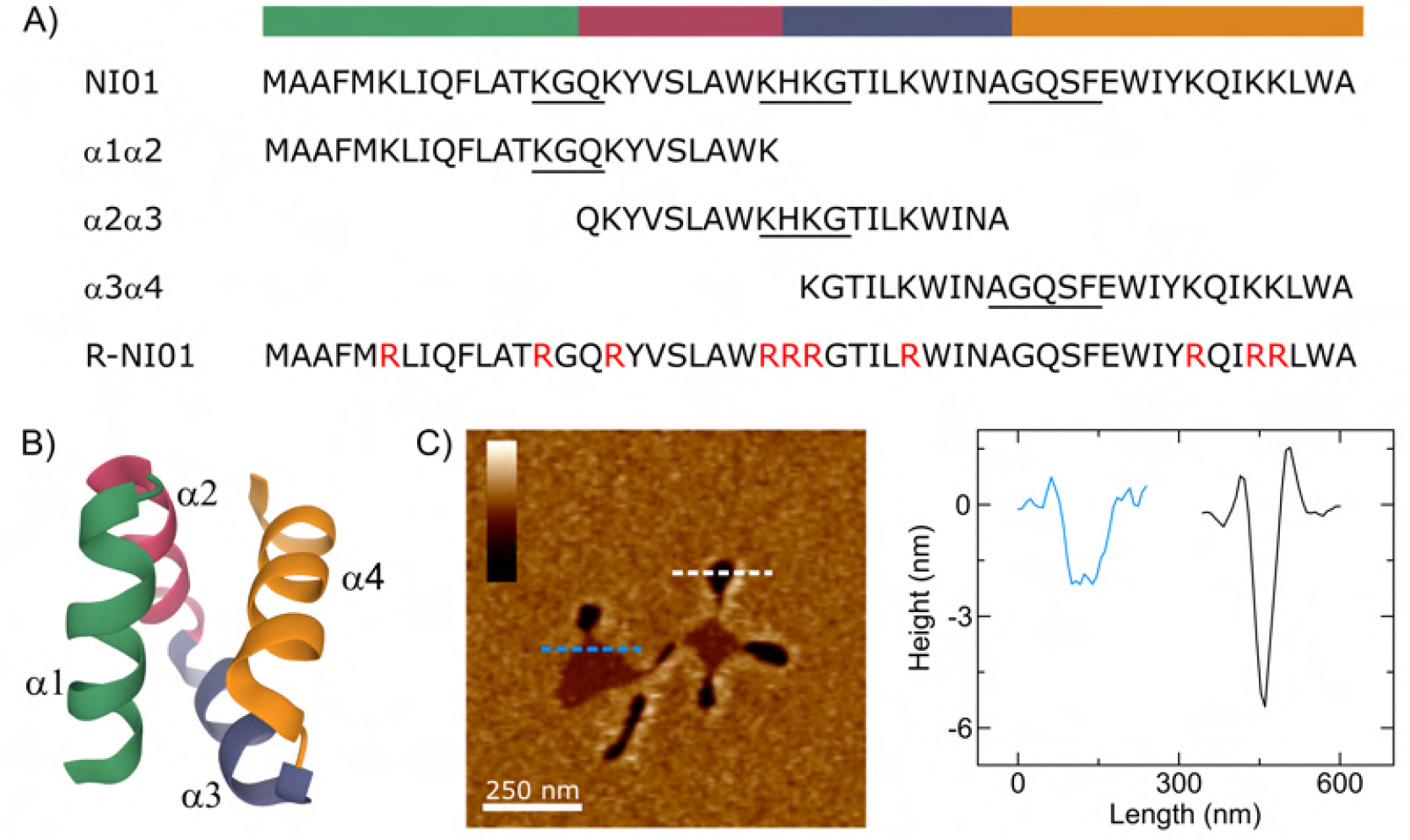
Overview of the NI01 peptide series and the observed dual mechanism against anionic lipid bilayers. ^9^ A) The sequences of epidermicin NI01 primary structure, its 2-helix substructure derivatives, and the arginine mutant R-NI01. The three substructures are notated as (*α*1*α*2, *α*2*α*3 and *α*3*α*4). The four helices are color coded above the sequences and labeled *α*1-*α*4 accordingly. Turns are underlined in the sequences. B) Crystal structure of NI01. Ribbon representation from the N terminus (green) to the C terminus (orange). C) In-liquid AFM images representing the topography of NI01-treated PC/PG (3:1) SLBs mimicking bacterial membranes. The floral shaped topography consists of bilayer spanning petal shaped pores (dark regions) and a membrane thinning patch in the center (light regions).

### NI01 mode of action

NI01 exhibits a synergistic multiple mechanism in anionic phospholipid bilayers characterized by the formation of transmembrane lesions and pores and membrane thinning patches, ~2 nm in depth half-way through the bilayer, in a floral pattern (Figure 1C).^9^ NI01 is also non-stereoselective as its all-D form demonstrated similar membrane disruption patterns. Given this mechanism, we sought to examine the modes of action for both the all-L and all-D forms of NI01 (Figure 2). Firstly, we found a strong dependence of GUV survival on peptide concentration - when treated with 5 μM of the peptide, the vast majority (~89.6%) of the GUVs survived treatment (i.e., no dye leakage was observed) over the timescales measured. However, for higher concentrations (10 μM and above), the entire GUV population showed complete leakage of the encapsulated dye molecules within the timescales of the experiment, with the mean time of leakage decreasing with increasing concentration. The large fraction of vesicle survival when dosed with 5 μM of the peptide may result from the peptide not reaching the threshold concentration necessary to induce leakage in individual vesicles. For the remaining peptide dosages, LEs converged into a unimodal narrow distribution around a concentration-specific timepoint (LEs histograms are depicted in the SI). The LEs’ mean time point inversely correlated with the corresponding peptide concentration (10 μM: 322.2 ± 44.0 min N=388, 25 μM: 136.2 ± 34.6 N=469, 50 μM: 74.4 ± 14.8 N=599). This indicates that NI01 is not a stochastic pore former in GUVs since its action is concentration dependent and the LEs are monodisperse. The heat maps for the all-D form experiments demonstrate almost identical response to the all-L form (Fig 2B). Therefore, both epimeric forms exhibit similar modes of action consistent with the mechanism established by AFM.

**Figure 2:**
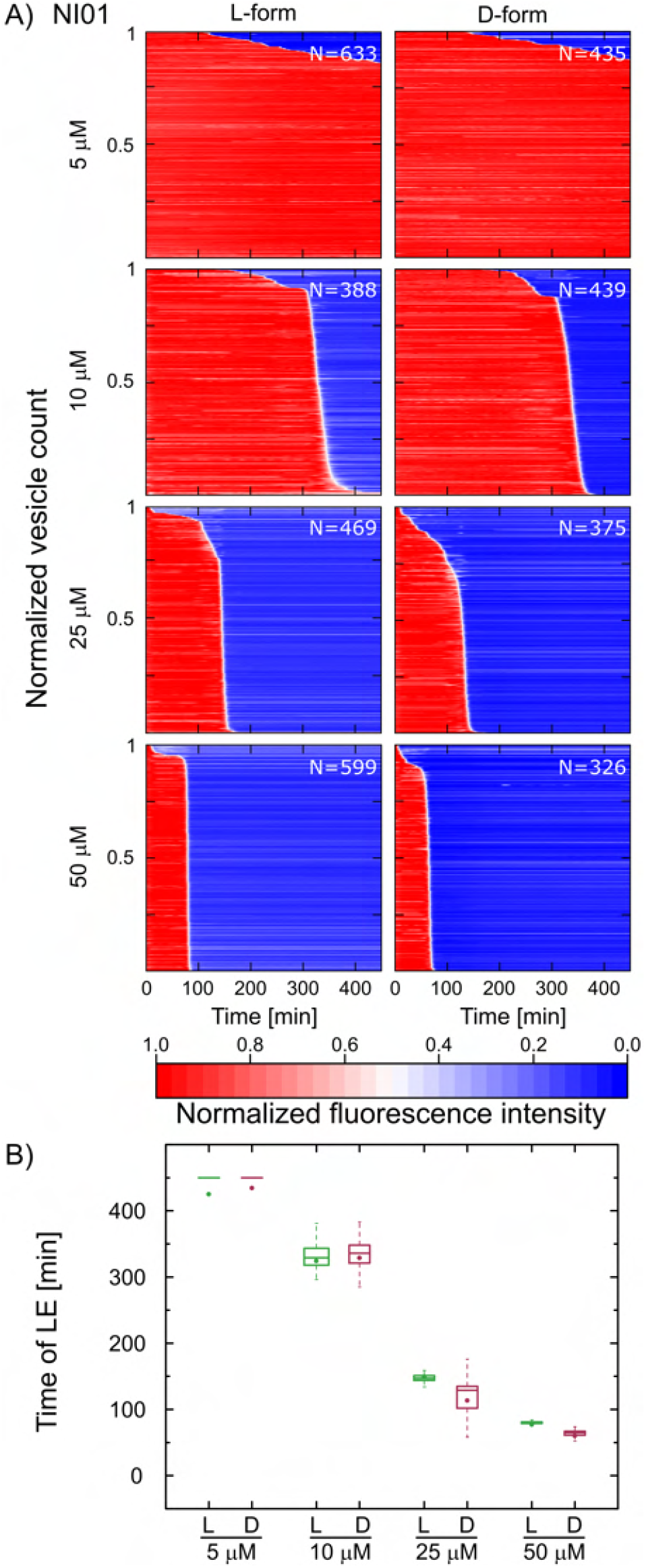
The all L- and all D- forms of NI01 compromise model bacterial membranes in a dose-dependent manner, showing similar modes of action. A) The heat plots map the mode of action by summarizing the leakage kinetics of NI01 treated populations of PC/PG (3:1) GUVs. Each horizontal line depicts the normalized intensity of encapsulated HPTS dye in a single trapped vesicle over time. The vesicle’s membrane is considered intact at high fluorescence intensity (red) and compromised at low fluorescent signal (blue). The intensity traces were ordered by the LE time point. Total number of GUVs is displayed per experiment. B) Box plot comparing the time distribution of LEs for both NI01 variants.

### Dissecting NI01 to understand the relation between its mechanism and mode of action

The observed dual mechanism of the four-helix bacteriocin was further examined by deconstructing the peptide into three 2-helix hairpin substructures and monitoring the modes of action for each (Figure 1A). Each of the hairpins supports a specific part of NI01’s multiple mechanisms:^9^ *α*1*α*2 and *α*2*α*3 hairpins form elongated transmembrane lesions and membrane thinning patches, respectively (figure S5). In contrast, *α*3*α*4 combines the formation of thinning patches with that of circular transmembrane pores (figure S5). We therefore probed whether the differences in mechanisms for the three hairpins translate to different modes of action. The experiments were conducted at 4 different concentrations for each substructure. LEs were observed at the 50 μM concentration. At the lower doses, most of the GUVs survived the treatment within the timescales measured. Compared to the data for NI01 (Fig 2), this indicates that the efficacy of the substructures against GUVs is lower than the activity of the full-size NI01. At the 50 μM concentration for the *α*1*α*2 and *α*3*α*4 hairpins, we observed the total leakage of all the individual vesicles. The timing of the LEs grouped into a unimodal narrow distribution around a similar timepoint (240.1 ± 10.7 min N=477 and 267.4 ± 10.8 min N=408) for both peptides, respectively. In regards to *α*2*α*3, the recorded timing of LEs showed some stochasticity and did not converge around a single time point. At the end of the experiment (450 min), the population was split into ~45% survivors and ~55% disrupted vesicles. These results highlight a correlation between the membrane active mechanism of a certain peptide with its mode of action as defined via a signature LE time distribution. Peptides that introduce transmembrane lesions have similar modes of action on populations of GUVs. With regards to the mechanisms of NI01 and *α*3*α*4, the mode of action appeared to be dictated by transmembrane poration as the dominant mechanism over membrane thinning. Similar to NI01, the substructures do not qualify as stochastic pore formers in GUVs since peptide action is concentration dependent and LEs only occur after a certain lag time.

### Exploring correlations between the mechanism and mode of action of NI01

Electrostatic interactions are one of the main driving forces responsible for the membrane activity of antimicrobial peptides. To gain a better insight into the role of charge on the activity of NI01, an all-arginine mutant of NI01 was synthesized, in which all cationic residues in the native sequence were replaced with arginines to ensure stronger electrostatic interactions with anionic membrane surfaces.^27^ This difference in membrane interaction is believed to arise from the fact that arginine is positively charged at all stages of membrane insertion, whereas lysine can become deprotonated once in the membrane. Similar to the other peptide, the mode of action for the mutant was probed using four different peptide concentrations (5, 10, 25 and 50 μM). A high percentage of LEs was observed even at the 5μM concentration indicating that the efficacy of the mutant against GUVs appeared higher than that of the original NI01 structure (Survival: NI01=89.6%, R-NI01= 0%). Furthermore, the distribution of timepoints of vesicle LEs at the lower peptide concentrations proved to be wide (5 μM: 122.6 ± 90.4 min N=440, 10 μM: 102.8 ± 76.2 min N=339, 25 μM: 134 ± 74.4 min N=479). Increasing the peptide concentration led to a second peak and the convergence of LEs clustering around a specific, concentration-dependent time point (50 μM: 103.0 ± 32.3 min N=518), (Fig. S3). This suggests that at 50 μM, a threshold concentration is reached. At this concentration LEs clustered into a unimodal narrow distribution at the mentioned timepoint. For R-NI01, thinning patches were the mechanism identified via AFM measurements, with no detection of the transmembrane pores that were observed with the original NI01 peptide (figure S5).^9^ Given this, it is interesting that, as mentioned above, the vesicle studies revealed strong membrane compromising activity even with 5 μM of the mutant peptide, in comparison to the original NI01 peptide at this concentration. The fact that R-NI01 has a strong effect on the GUVs is, however, consistent with the stronger peptidelipid interactions furnished by arginine residues. Unlike lysine, arginine provides five hydrogen-bond donors forming stable peptide–phosphate clusters, which enhance the affinity of the peptide to phospholipids changing the dynamics of membrane disruption to favour membrane exfoliation.^27,28^ However, comparing the time distributions of LEs between NI01 and R-NI01 (Fig 4B), we can say unequivocally that, as far as the mode of action is concerned, R-NI01 qualifies as a stochastic pore former in GUVs since LEs occur without any specific lag period and are spread over a wide range of time points (Figure S3).

**Figure 3:**
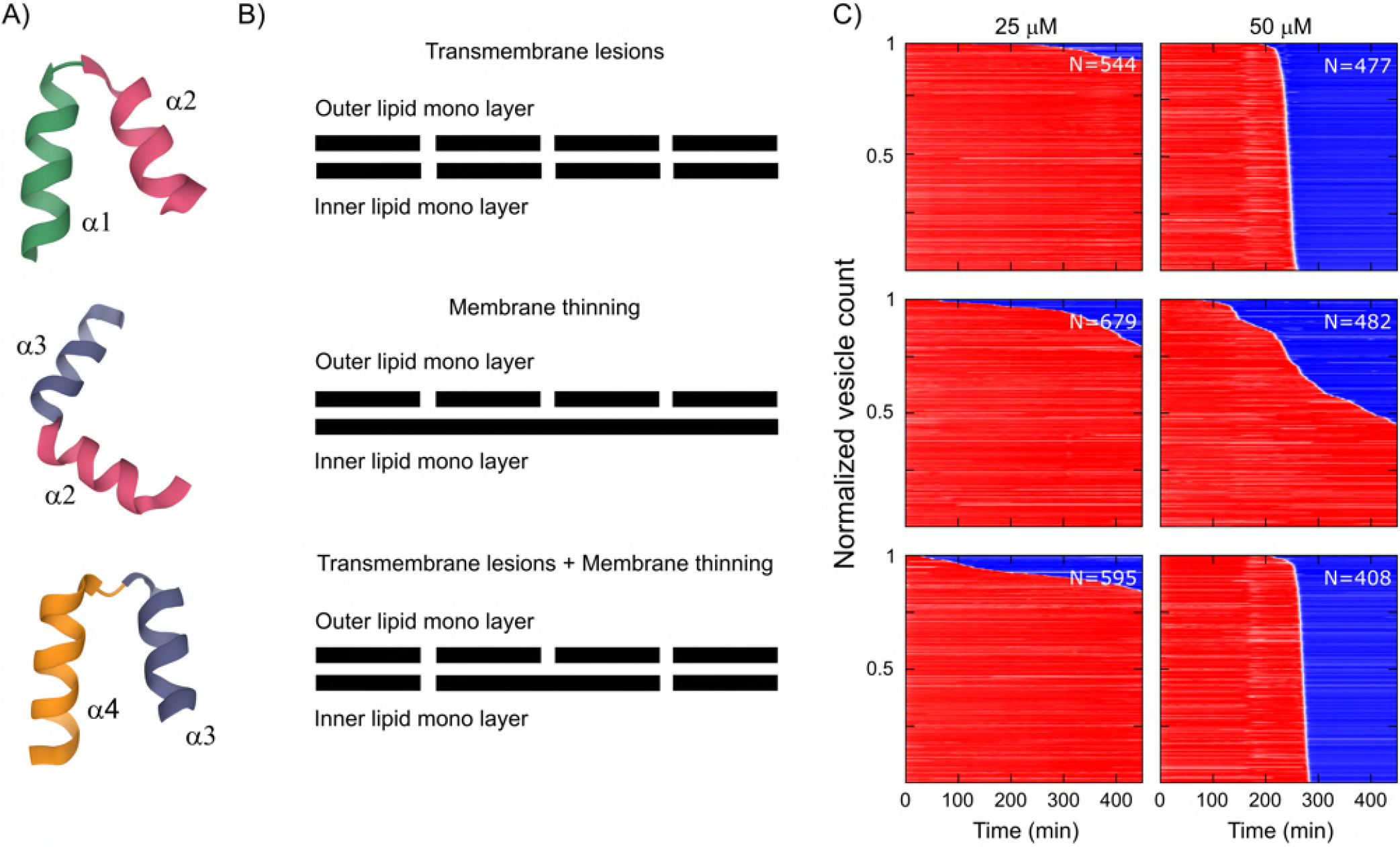
Dissecting multiple mechanisms of NI01 into substructure derivatives and corresponding modes of action. A) Three 2-helix substructures derived from NI01. B) Schematics representing the different mechanisms for each of the substructures as interpreted from the in-liquid AFM topography images of anionic PC/PG (3:1) SLBs treated with the two-helix hairpins (AFM images are provided in the SI) C) Heat plots mapping the modes of action of the three sub-structures, *α*1*α*2, *α*2*α*3 and *α*3*α*4 against anionic PC/PG (3:1) GUVs.

**Figure 4:**
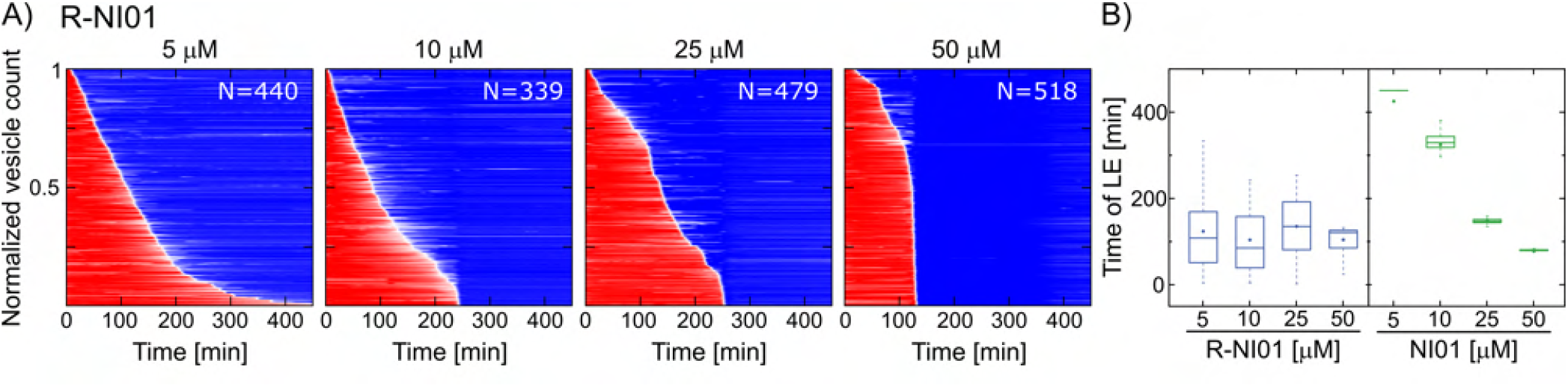
Exchanging lysines with arginines in NI01 transforms its mode of action. A) Heat plots mapping the mode of action of the mutant R-NI01 against anionic PC/PG (3:1) GUVs. B) Box plot comparing the LE time distributions between the native NI01 and mutant R-NI01 at 4 different concentrations.

### Exploring model mechanisms and correlating modes of action

To further explore correlations between mechanisms of membrane disruption and modes of action, we investigated a series of archetypal and well-studied AMPs. These were chosen to allow for a comparison to the inferences drawn from the NI01 series. These peptides have different molecular mechanisms and modes of action. Melittin and magainin 2 are known to form toroidal pores with the distinction of having graded and all-or-none leakage characteristics, respectively.^29–31^ Graded action refers to the mode where all the individual vesicles show similar leakage characteristics, and hence the time points of LEs are tightly distributed (unimodal). On the other hand, all-or-none action indicates that the population can be split into intact and leaked vesicles (bimodal). Alamethicin is postulated as a model peptide for barrel-stave pore formation.^32,33^ Barrel-stave and toroidal pores are both transmembrane. However, the walls of barrel-stave pores only consist of peptides, while toroidal pores cause the membrane to bend, with the phospholipid headgroups facing the interior of the pore. Polymyxin B lyses membranes in a detergent-like manner, ^24^ while the mechanism of the linear form of hβD-3 is believed to involve disordered toroidal pore formation.^34^ The concentration at which these peptides were studied, the lipid composition of reconstituted membranes used in the studies, and other experimental parameters are factors that may affect their activity characteristics. With this in mind, we characterized the modes of action of these peptides using the GUV studio under the conditions used for the first series, to probe unique signatures for the LE time distributions for these peptides. We then correlate the LE time-distribution with the modes of action of the peptide against their molecular mechanisms.

First, we compared the LE time distributions of melittin and magainin 2. For melittin, we detected the total leakage of the entire vesicle population at all concentrations but 5 μM - a scenario reminiscent of NI01’s mode of action. For magainin 2, LEs corresponding to most of the vesicle population were observed only at 25 and 50 μM. Moreover, at 10 μM the peptide displayed a unique LE time distribution where a ~44% subpopulation of vesicles remained intact throughout peptide treatment. With the process of vesicle leakage being stochastic, the time distribution of LEs that took place converged around a certain time point (10 μM: 116.1 ± 77 min N=177) suggesting that magainin 2 forms transmembrane pores, which is in contrast to the effects observed for *α*2*α*3 characterized by the mechanism of membrane thinning. These results concur with those reported by others for melittin and magainin 2.^29–31^ Although both peptides by virtue of forming transmembrane pores exhibit a similar molecular mechanism, our results suggest that their *modes of action* are different in that melittin shows a graded characteristic, while magainin 2 favours the all-or-none characteristic.^35^

We then compared melittin with alamethicin. Both peptides gave comparable LE time distributions at different concentrations, similar to what was observed for NI01. The majority of LEs occur after a certain lag time (10 μM: 165.3 ± 62.6 min N=406, 25 μM: 61 ± 17.9 min N=641, 50 μM: 34.1 ± 23.8 min N=645), which inversely correlated with corresponding peptide concentrations, thus indicating that both the toroidal and barrel-stave poration follow the two-state model.^13,36^ Next, our assay revealed that polymyxin B was the most potent amongst all of the tested peptides. The total leakage of all the vesicles in the population was observed at all concentrations and within shorter timescales than for the other peptides. The LEs converged around a certain time point (5 μM: 92.9 ± 33 min N=505, 10 μM: 47.1 ± 34.94 min N=388, 25 μM: 28.6 ± 35.9 min N=480, 50 μM: 11.5 ± 16.9 min N=430) supporting the previous two-state model, as with increasing peptide concentration the lag time duration before LEs reduced.

Lastly, the linear form of hβD-3 demonstrated a mode of action analogous to that of R-NI01. The total leakage of >~80% of the vesicle population was observed at each concentration used, and in all cases the time distribution for LEs was wide, skewed and qualitatively similar (5 μM: 182.3 ± 153.4 min N=569, 10 μM: 179.9 ± 164.4 min N=524, 25 μM: 123.6 ± 153 min N=324, 50 μM: 59.9 ± 82.6 min N=589). Increasing the peptide concentration resulted in narrowing the skewed distribution by decreasing the lag time to the LE while increasing the duration of vesicle leakage (i.e., the amount of time for the dye to leak out of an individual vesicle). Since the LEs do not occur after a specific lag period, the latter observation qualifies the peptide as a stochastic pore formation agent in GUVs.

## Conclusion

Typically, a molecular mechanism is assigned to an antimicrobial peptide to describe the nature of its interactions with phospholipid bilayers. Assigning a mode of action, which reflects the kinetics of peptide-membrane interactions and may provide correlations with the underlying molecular mechanisms, is less common. We designed and applied a bespoke GUV studio to determine the modes of action of a range of antimicrobial peptides. The study provided important insights into correlations between the molecular mechanisms of the peptides and their modes of action. Firstly, for the epimeric forms of a peptide demonstrating the same molecular mechanism, the mode of action is also the same. Secondly, if a peptide exhibits multiple mechanisms, the main determinant of its mode of action is its most disruptive mechanism. Thirdly, LEs caused by transmembrane poration follow one of two modes of action. In one, a two-state mode, pore formation ensues upon reaching a threshold peptide concentration. This mode is characterized by LEs occurring within a narrow time range, with peptide concentration determining the mean time point of the LEs. Another one is a spontaneous mode, where LEs can take place before reaching a threshold peptide concentration. This leads to LEs that are distributed over a wide range of time points, for which the impact of peptide concentration is limited to the width of the LE time distribution.

Fourthly, pore forming peptides demonstrating different molecular mechanisms, e.g. melittin and alamethicin, can share a similar mode of action. By contrast, peptides exhibiting the same molecular mechanism, e.g. melittin and magainin 2, can demonstrate different, e.g. “graded” or “all-or-none”, modes of action. Finally, GUV responses to peptides inducing membrane thinning typically correlate with a mode of action that is associated with stochastic LEs. The majority of the vesicle population in this scenario remains intact. The demonstrated correlations between the molecular mechanisms of peptides and their kinetic modes of action suggest that the sole use of MIC measurements to define antimicrobial treatments requires reconsideration. The direct comparisons allowed by the GUV studio offer an effective approach to systematically characterize membrane-active antimicrobials. By expanding this methodology to a larger number of peptides representing different antimicrobial mechanisms, it will be possible to systematically characterize their different modes of action in a standardized manner, which may better inform the design and translation of clinically acceptable peptides.

## Experimental Section

### Optofluidic leakage Assay

The optofluidic GUV leakage assay builds on our previous work studying the activity of Cecropin B.^19^ The assay utilizes the GUV studio to standardize experimental conditions and overcome numerous technical drawbacks that classical leakage techniques suffer from. The assay is designed to monitor the leakage kinetics of the encapsulated fluorophore Pyranine from a population of GUVs at single vesicle resolution, under continuous peptide administration. Our microfluidics platform facilitates both control over the dosed peptide concentrations and the imaging of hundreds to thousands of GUVs at desirable spatiotemporal resolution. Importantly, the continuous perfusion of the peptides ensures that all the GUVs studied are exposed to similar amounts of the peptides simultaneously, negating any potential innoculum effect.^37^

### GUV studio

As described in Figure 6, the GUV studio incorporates an on-chip GUV formation component that relies on the octanol-assisted liposome assembly technique.^38^ This is followed by a downstream vesicle immobilization component consisting of 8 separate chambers each containing an array of 720 vesicle traps aimed to capture vesicles with diameters in the range of 20-25 μm. The connector channel (shown in yellow) integrates the above mentioned components and its main purpose is to efficiently distribute the vesicles generated into the 8 chambers. Each chamber is connected to a perfusion inlet responsible for peptide administration and enables the total exchange of fluid surrounding the GUVs in a highly controlled manner. Therefore, eight independent experiments can be run in parallel using a single device. The GUV studio is equipped with a pneumatic valve system consisting of 3 control channels (shown in red) that offer enhanced control when switching between the vesicle formation and trapping stage and the peptide administration stage. The microfluidic flows for all depicted inlets were controlled using the Fluigent MFCS-EZ flow control system (Fluigent S.A, France) and its accompanying MAESFLOW software. Inlets were connected to the fluid reservoirs (Micrewtube 0.5 mL, Simport) via a polymer tubing (Tygon microbore tubing, 0.02” × 0.06”, Cole Parmer, UK) and a metal connector tip (Gauge 23 blunt end, Intertronics). A neMESYS syringe pump with a 250μL Duran Borosilicate glass syringe (ILS, Germany) was connected to outlet A via Upchurch 1520 G tubing (0.03” × 0.06”) (figure 6). A more detailed description of the device operation can be found in the SI.

**Figure 5:**
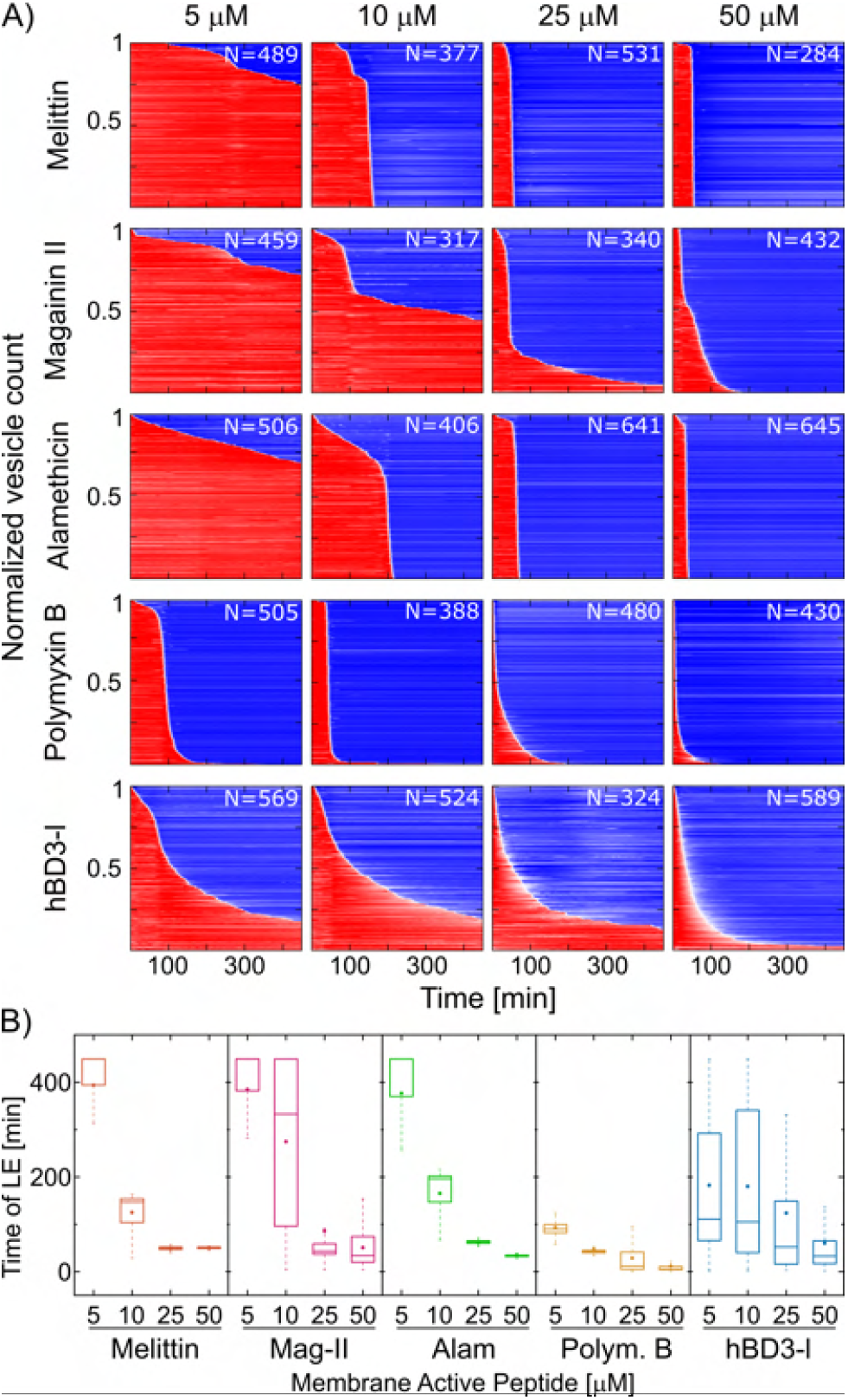
Investigating the mode of action of archetypal AMPs exhibiting various molecular mechanisms. A) Heat plots mapping the disruption activity and mode of action of melittin, magainin 2, alamethicin, polymyxin B and hβD-3-l in anionic PC/PG (3:1) GUVs. B) Box plot comparing the time distribution of LEs for the mapped series.

**Figure 6:**
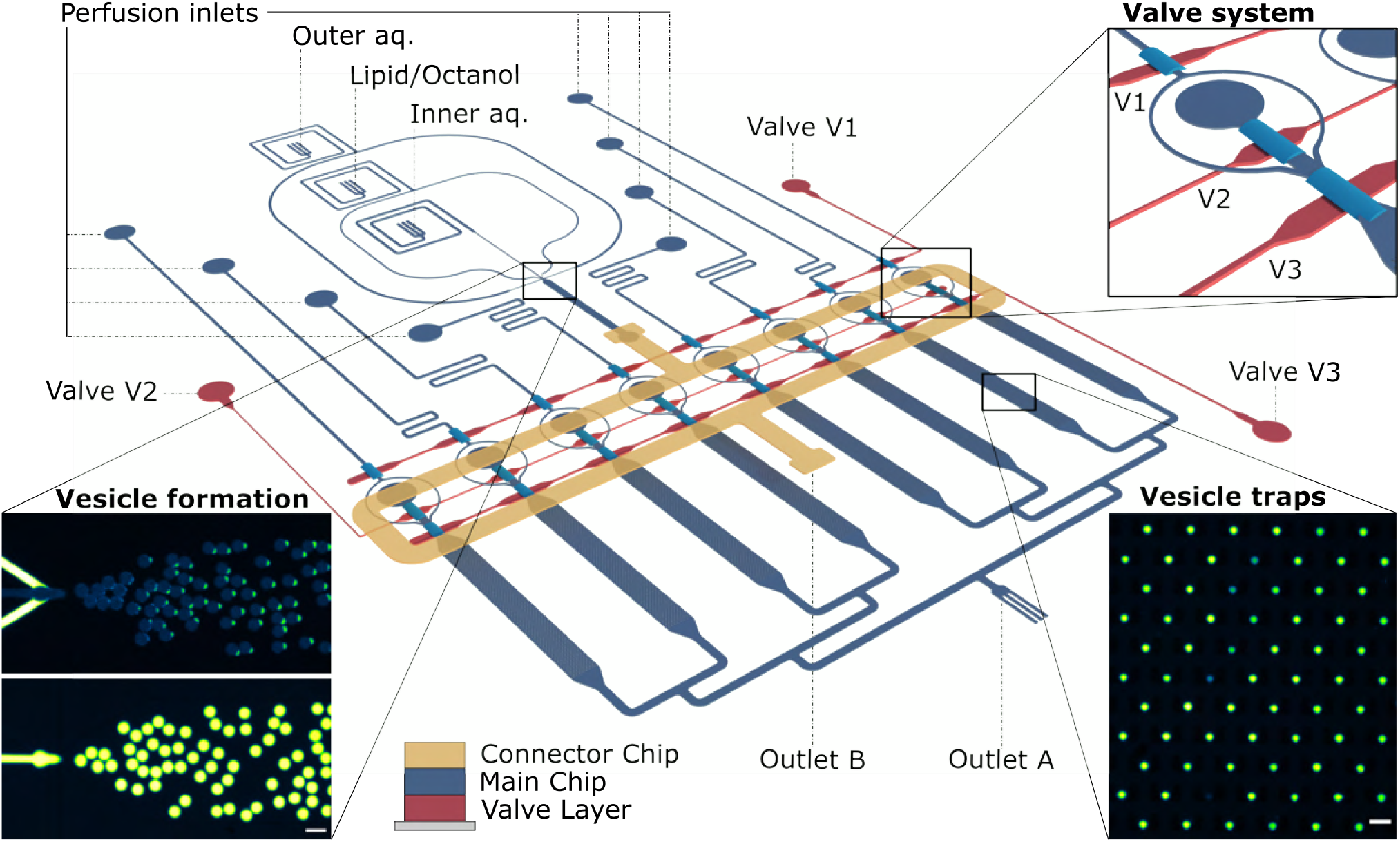
Overview of the GUV sutdio, a second-generation microfluidic platform utilized for characterising the discussed membrane-active peptides. ^19^ The schematic outlines the platform’s capabilities in preparing GUVs, vesicle trapping, and controlled peptide adminstration. The GUV formation element is based on the octanol-assisted liposome assembly (OLA) technique. The platform integrates 8 separate chambers each encompassing an array of 720 vesicle traps. Vesicles are transported from the formation junction to the trap arrays via a second layer of microfluidic channels, labelled as the connector chip (in yellow). This design features a pneumatic valve system in a third layer underneath the main chip (in red). The valve system consists of 3 control channels that can be used to facilitate controlled switching between the unique stages of the experiment. Insets show microscopic fluorescence images of vesicle formation and trapping. The lipid phase is tagged with Liss Rhod PE, the GUVs are encapsulating HPTS dye. Scale bars represent 50 μm.

### Data acquisition and analysis

The device was mounted on an Olympus IX73 inverted microscope equipped with an automated Prior XYZ-stage, a Photometrics Evolve 512 camera coupled with a wLS LED lamp (QImaging), a FITC filter cube set (Chroma) and a 10× air objective (Olympus UPLFLN). The software μManager 1.4 was used to synchronize the camera acquisition and the movement of the automated stage. Each chamber was designed such that it could be imaged within 8 fields of view (FOVs), hence an image acquisition cycle over the entire device consisted of 64 FOVs. The automated stage takes 1 min to scan all 64 FOVs. Vesicle monitoring continued overnight and we reported 450 cycles after peptide arrival in the GUV trap chambers. The acquired videos were analysed using a bespoke automated python code to detect GUVs and obtain fluorescence signal traces for individual vesicles (SI for further details).

### Microfluidic Chip Fabrication

The GUV studio is a multilayered polydimethylsiloxane (PDMS) device that relies on standard photolithography and soft-lithography techniques.^39^ Three molds were fabricated matching the valve control layer (bottom), main chip (middle) and connector chip (top) as depicted in (fig.6). The main chip channels were molded from a master with two distinct channel profiles, (1) rectangular features with heights of ~20 μm and ~30 μm, and (2) rounded features with heights of ~25 μm. The mold of the connector channel has features of ~35 μm in height, whereas the features of the valve control layer are ~30 μm in height. The SU-8 2025 negative photoresist was used for rectangular features with heights around 20, 30 and 35 μmby spin-coating at 3800, 2800, and 2300 rpm respectively for 60 s with a ramp of 100 rpm s^*−*1^ on 4” silicon wafers (University Wafer, USA). The round channels required spin-coating AZ 40XT-11D positive photoresist at 2500 rpm on top of the SU-8 features. The spin coating was followed by a soft bake step where in the case of the negative resist, the coated wafer was placed on a hot plate for 1 min at 65°C and then 5 min at 95°C. In the case of the positive resist, the temperature was gradually increased from 80°C to 125°C over 7 min. A high-resolution, table-top laser direct imaging system (LPKF ProtoLaser LDI, Germany) was used for prototyping on the resist-coated substrates directly using AutoCAD designs. The mold then was post baked for 1 min at 65°C followed by 5 min at 95°C in the case of the negative resist, and 2 min at 100°C for the positive resist. The devices were developed in PGMEA for 3 min (SU-8) or AZ 726 MIF (AZ). Then the negative resists molds were hard-baked for 15 min at 125°C, and the positive resist mold was baked for ~7 min at 120°C to re-flow the resist and obtain the round channels.^39^ To achieve different feature types and heights on the same silicon wafer, the photolithography process was performed three times on the same silicon wafer using a built-in anchoring tool for aligning the features.

PDMS replicas of the connector and main chips were obtained from the Silicon mold by mixing the elastomer with the curing agent (Sylgard 184, Dowsil) in a 9:1 ratio. The PDMS mixture was then degassed for 30 min in a desiccator, poured onto the molds and baked in the oven for 90 min at 60°C. The valve control PDMS membrane was obtained using a mixing ratio of 14:1 (elastomer:curing agent) and spincoating the PDMS on the mold at 1500 rpm for 60 s with a ramp of 100 rpm s^*−*1^. After curing and peeling off the main chip, the inlets/outlet were punched with a 0.75/1.5 mm biopsy punch (WPI, UK) respectively. A glass slide (76 mm × 38 mm × 1 mm) was coated with PDMS and used as the base for the devices. The main chip was aligned and bonded to the PDMS valve control membrane by exposing the surfaces to an oxygen plasma (100 W plasma power, 10 s exposure, 25 sccm, plasma etcher from Diener Electric GmbH & Co. KG) and then annealing the exposed surfaces. After bonding, the main chip was peeled-off and the valve control inlets created using the 0.75 mm biopsy punch. The sample was then bonded to the PDMS-coated glass slide using oxygen plasma. The surface of the microfluidic channels delivering the outer aqueous solution (OA) was treated for 15 min with polyvinyl alcohol (PVA) solution (50 mg mL^*−*1^, 87-90% hydrolysed, Sigma-Aldrich).^19,38^ Post treatment, the PVA was removed via vacuum pump suction and baked at 120°C for 15 min. Finally, the connector PDMS layer was aligned and plasma bonded on top of the main chip.

### GUV preparation

The Inner Aqueous (IA) solution of GUVs consisted of a PBS buffer (10 mM phosphate buffer, 2.7 mM KCl and 137 mM NaCl) at pH 7.4, in addition to 200 mM sucrose and 15% v/v glycerol. The Outer Aqueous (OA) solution additionally contained 50 mg mL^*−*1^ of Kolliphor P-188 (Sigma-Aldrich, UK) as per standard OLA protocols.^38^ Lipids were purchased from Sigma Aldrich in powder form and were dissolved in 100% ethanol to a final concentration of 100 mg mL^*−*1^. The lipid/octanol phase (LO) was prepared by diluting the lipid stock in 1-octanol to a 4-5 mg mL^*−*1^ concentration. A lipid mixture of 3:1 ratio 1,2-dioleoyl-sn-glycero-3-phosphocholine (DOPC) with 1,2-dioleoyl-snglycero-3-phospho-rac-(1-glycerol) sodium salt (DOPG) was used to mimic the anionic charge of typical bacterial membranes. ^19^ 16:0 Liss Rhod PE (Avanti Lipids, 0.5 mg mL^*−*1^ in ethanol) was doped to label the LO phase. To monitor dye leakage from GUVs, HPTS (8-hydroxypyrene-1,3,6-trisulfonic acid, Thermo Fisher) was diluted into the inner aqueous phase (50 μM).

### Peptide preparation

NI01 and all its derivatives were synthesized, identified and purified as previously published.^9^ Briefly they were assembled in a Liberty microwave peptide synthesizer (CEM Corp.). All peptides were then purified by semi-preparative RP-HPLC. The purity and identities of NI01 and derivatives were confirmed by analytical RP-HPLC (≥ 95%) and MALDI-ToF mass-spectrometry. Analytical and semi-preparative RP-HPLC was performed on a Thermo Scientific Dionex HPLC System (Ultimate 3000) using Vydac C18 analytical and semi-preparative (both 5 μm) columns. Analytical runs used a 10-70% B gradient over 30 min at 1 mL/min, semi-preparative runs were optimized for each peptide, at 4.5 mL min^*−*1^. Detection was at 280 and 214 nm. Buffer A and buffer B were 5% and 95% (v/v) aqueous CH3CN containing 0.1% TFA. Melittin, Magainin 2, Alamethicin (all at HPLC purity ≥97%) and Polymyxin B Sulfate were purchased from Merck, UK. The linear hβD-3 variant was synthesized and purified as previously described.^34^ The lyophilized peptides to be tested were freshly hydrated in Milli-Q water briefly before the experiment except for Alamethicin due to its poor solubility in water, the Alamethicin stock was dissolved in Ethanol. Aliquots were then diluted to the desired peptide concentrations in the same solution as the IA phase; a small amount of HPTS (5 μM) was also added as a tracer to precisely time peptide arrival in the GUV trap chambers.

### In-liquid AFM imaging of SLBs

AFM experiments were conducted as described in detail previously.^9^ Briefly, small unilamellar vesicles (SUVs) were prepared using lipid film hydration and extrusion protocols. A 3:1 lipid film of 1-palmitoyl-2-oleoyl-glycero-phosphocholine (POPC) with 1-palmitoyl-2-oleoyl-sn-glycero-3-phospho-(1’-rac-glycerol) (POPG) lipids, Avanti Polar Lipids (Alabaster, USA) was hydrated and sonicated with 10 mM phosphate buffer (pH 7.4). The solution was then extruded through a polycarbonate filter (0.05 μm) using a hand-held extruder (Avanti Polar lipids) to a final concentration of 1 mg mL^*−*1^. To form SLBs on mica, SUVs were incubated for 45 min onto cleaved mica (Agar Scientific, UK) that was pre-hydrated in imaging buffer (20 mM MOPS, 120 mM NaCl, 20 mM MgCl2 (pH 7.4)). The sample was then washed 10 times with imaging buffer. The topographic imaging of SLBs in aqueous buffers was performed on a Multimode 8 AFM system (Bruker AXS, USA) using Peak Force Tapping™ mode and MSNL-E cantilevers (Bruker AFM probes, USA). Images were taken at a PeakForce frequency of 2 kHz, amplitude of 10-20 nm and set-point of 10-30 mV (<100 pN). NI01 or its derivatives were introduced into a 100 μL fluid cell (Bruker AXS, USA) to the intended final concentration.

## Acknowledgement

The authors thank Rainer Bartels and Volker Grote for synthesis of the hβD-3 peptide. U.F.K. acknowledges support from an ERC consolidator grant (Designer- Pores 647144). K.A.N. acknowledges support from a Cambridge-National Physical Laboratory (UK) studentship, the Winton Programme for the Physics of Sustainability, the Trinity-Henry Barlow Scholarship and the ERC. M.F. acknowledges support from an EPSRC iCASE studentship. C.N. acknowledges funding from Phospholipid Research Center Heidelberg (Germany). J.C. acknowledges funding from a Wellcome Trust Institutional Strategic Support Award to the University of Exeter (204909/Z/16/Z). M.G.R. acknowledges funding from the UK’s Department for Business, Energy and Industrial Strategy and Innovate UK (103358). This research was funded in whole, or in part, by the Wellcome Trust (Grant number 204909/Z/16/Z). For the purpose of Open Access, the author has applied a CC BY public copyright license to any Author Accepted Manuscript version arising from this submission.

