## Supplementary Material for "Measuring thousands of single vesicle leakage events reveals the mode of action of antimicrobial peptides"

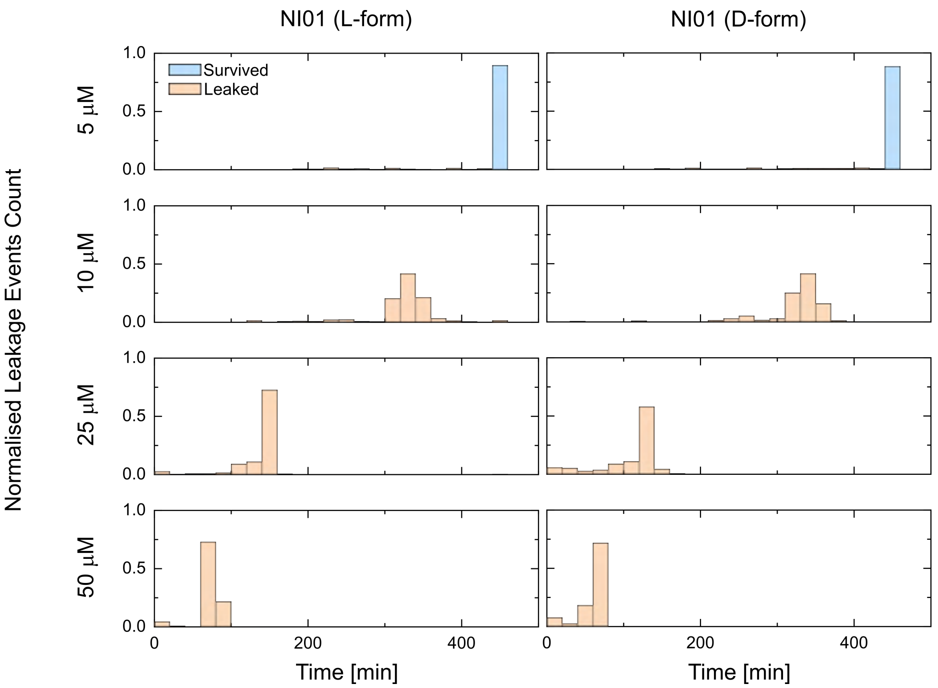


Figure S1: Side-by-side comparison between the all L- and all D- forms of NI01 using histograms depicting the LEs time distributions normalized to the count of GUVs monitored. The histograms correspond to the heat maps for Figure 2 in the manuscript.

Table S1: Summary of leakage events mean time point and GUV population survival within 7.5 hours of peptide treatment with the all L- and all D- forms of NI01.

| Peptide | Conc. | Average time | Std | Survival % |
|---|---|---|---|---|
| Epi N | 5 | 433.7 | 51.4 | 89.6 |
| Epi N | 10 | 322.2 | 44.0 | 0 |
| Epi N | 25 | 136.2 | 34.6 | 0 |
| Epi N | 50 | 74.4 | 14.8 | 0 |
| Epi N-D | 5 | 432.4 | 54.8 | 88.5 |
| Epi N-D | 10 | 320.8 | 48.1 | 0 |
| Epi N-D | 25 | 109.2 | 37.9 | 0 |
| Epi N-D | 50 | 57.6 | 15.7 | 0 |

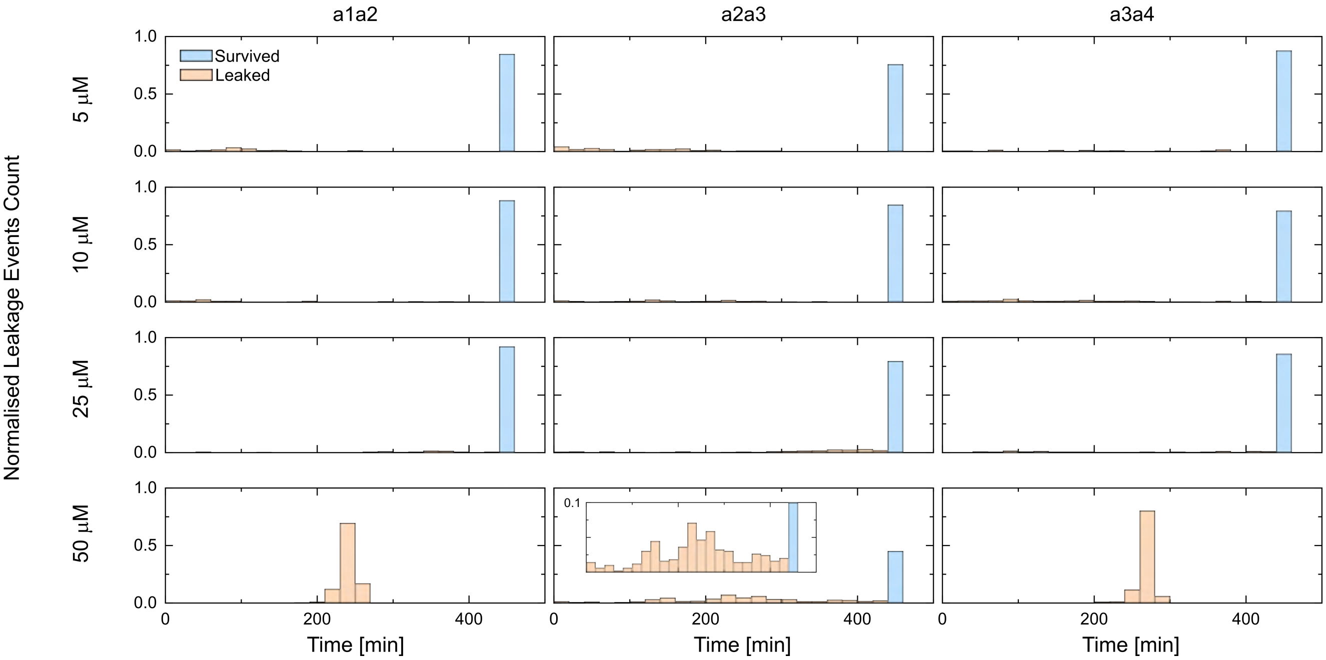


Figure S2: Side-by-side comparison between the three substructures of NI01 using histograms depicting the LEs time distributions normalized to the count of GUVs monitored. A subset was added for $\alpha 2\alpha 3$ at 50 µM to better visualize the wide distribution of LEs below 10% of the normalized count. The histograms correspond to the heat maps for Figure 3 in the manuscript.

Table S2: Summary of leakage events mean time point and GUV population survival within 7.5 hours of peptide treatment for the three substructures of NI01.

| Peptide | Conc. | Average time | Std | Survival % |
|---|---|---|---|---|
| $\alpha 1\alpha 2$ | 5 | 396.7 | 126.4 | 84.6 |
| $\alpha 1\alpha 2$ | 10 | 415.5 | 104.5 | 88.4 |
| $\alpha 1\alpha 2$ | 25 | 437.4 | 49.4 | 92.2 |
| $\alpha 1\alpha 2$ | 50 | 240.1 | 10.7 | 0 |
| $\alpha 2\alpha 3$ | 5 | 368.0 | 150.1 | 75.8 |
| $\alpha 2\alpha 3$ | 10 | 405.3 | 109.1 | 84.6 |
| $\alpha 2\alpha 3$ | 25 | 419.5 | 79.9 | 79.3 |
| $\alpha 2\alpha 3$ | 50 | 335.3 | 126.2 | 44.9 |
| $\alpha 3\alpha 4$ | 5 | 390.5 | 125.1 | 87.7 |
| $\alpha 3\alpha 4$ | 10 | 419.5 | 89.7 | 79.3 |
| $\alpha 3\alpha 4$ | 25 | 417.9 | 90.5 | 85.7 |
| $\alpha 3\alpha 4$ | 50 | 267.4 | 10.8 | 0 |

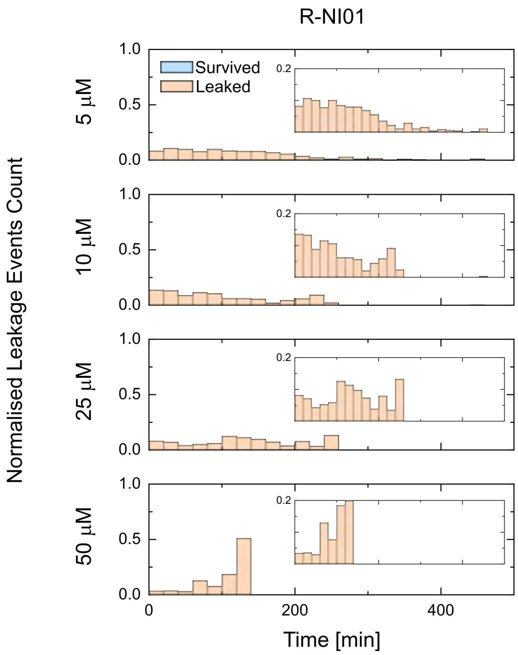


Figure S3: The time distribution of LEs induced by the mutant R-NI01 depicted in histograms normalized to the count of GUVs monitored. A subset was added for each concentration to better visualize the wide distribution of LEs below 20% of the normalized count. The histograms correspond to the heat maps for Figure 4 in the manuscript.

Table S3: Summary of leakage events mean time point and GUV population survival within 7.5 hours of peptide treatment with the mutant R-NI01.

| Peptide | Conc. | Average time | Std | Survival % |
|---|---|---|---|---|
| Epi R | 5 | 122.6 | 90.4 | 0 |
| Epi R | 10 | 102.8 | 76.2 | 0 |
| Epi R | 25 | 134.0 | 74.4 | 0 |
| Epi R | 50 | 103.0 | 32.3 | 0 |

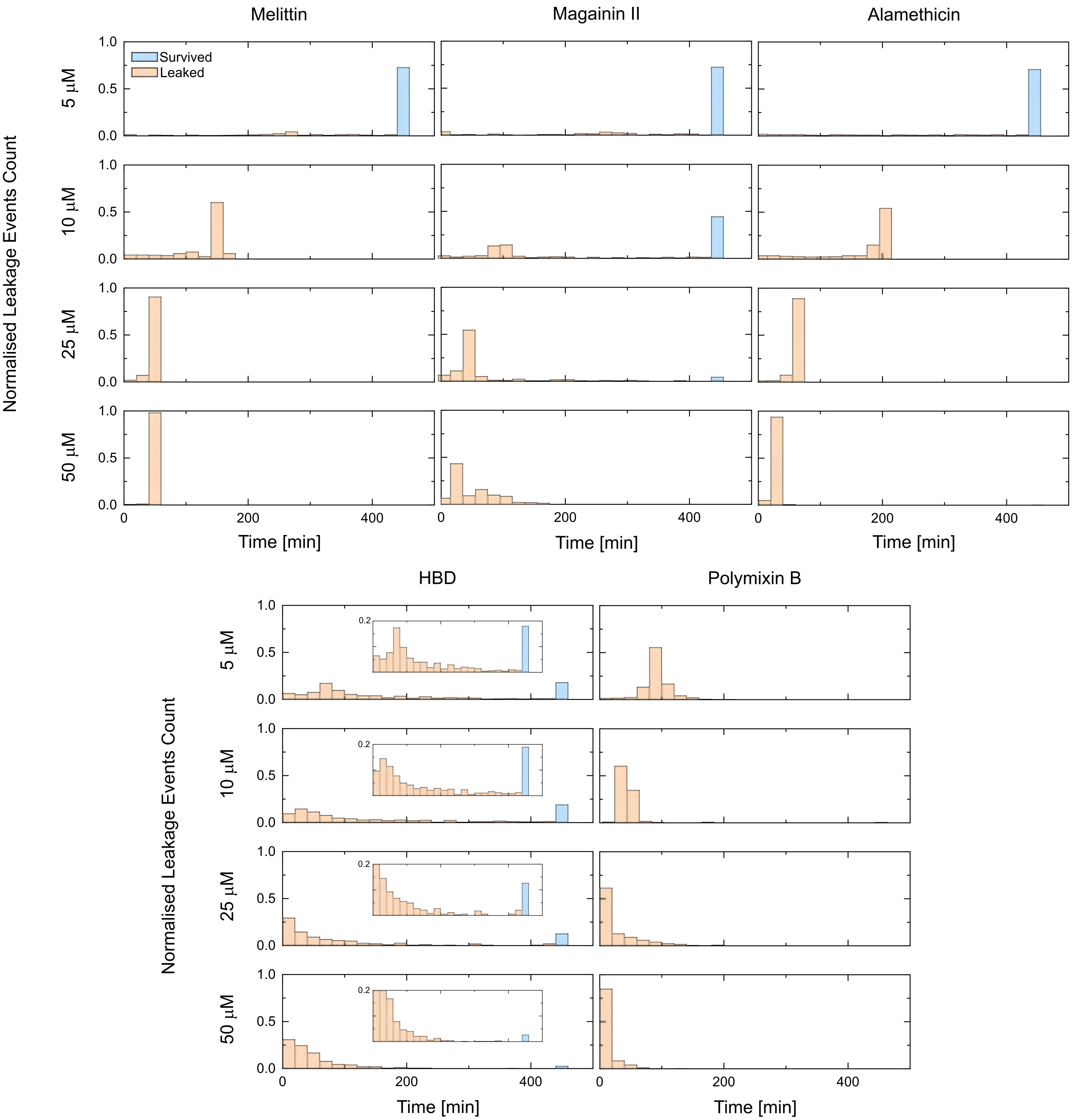


Figure S4: Side-by-side comparison between 5 archetypal AMPs of melittin, magainin 2, alamethicin, polymyxin B and hBD-3-l in anionic PC/PG (3:1) GUVs. Histograms depicting the LEs time distributions normalized to the count of GUVs monitored. A subset was added for each concentration for HβD-3 to better visualize the wide distribution of LEs below 20% of the normalized count. The histograms correspond to the heat maps for Figure 5 in the manuscript.

Table S4: Summary of leakage events mean time point and GUV population survival within 7.5 hours of peptide treatment with the 5 archetypal AMPs (melittin, magainin 2, alamethicin, polymyxin B and hBD-3-l).

| Peptide | Conc. | Average time | Std | Survival % |
|---|---|---|---|---|
| Melittin | 5 | 393.9 | 107.7 | 72.9 |
| Melittin | 10 | 124.9 | 44.9 | 0 |
| Melittin | 25 | 47.8 | 7.7 | 0 |
| Melittin | 50 | 50.0 | 4.4 | 0 |
| Mag-2 | 5 | 384.5 | 124.1 | 72.3 |
| Mag-2 | 10 | 274.7 | 173.9 | 44.3 |
| Mag-2 | 25 | 85.8 | 108.2 | 4.5 |
| Mag-2 | 50 | 50.7 | 37.6 | 0 |
| Alam | 5 | 376.6 | 133.9 | 70.9 |
| Alam | 10 | 165.3 | 62.6 | 0 |
| Alam | 25 | 61.0 | 17.9 | 0 |
| Alam | 50 | 34.1 | 23.8 | 0 |
| Polym | 5 | 92.9 | 33.0 | 0 |
| Polym | 10 | 47.1 | 34.9 | 0 |
| Polym | 25 | 28.6 | 35.9 | 0 |
| Polym | 50 | 11.5 | 16.9 | 0 |
| HBD | 5 | 182.3 | 153.4 | 18 |
| HBD | 10 | 179.9 | 164.4 | 18.9 |
| HBD | 25 | 123.6 | 153.0 | 12.6 |
| HBD | 50 | 59.9 | 82.6 | 2.69 |

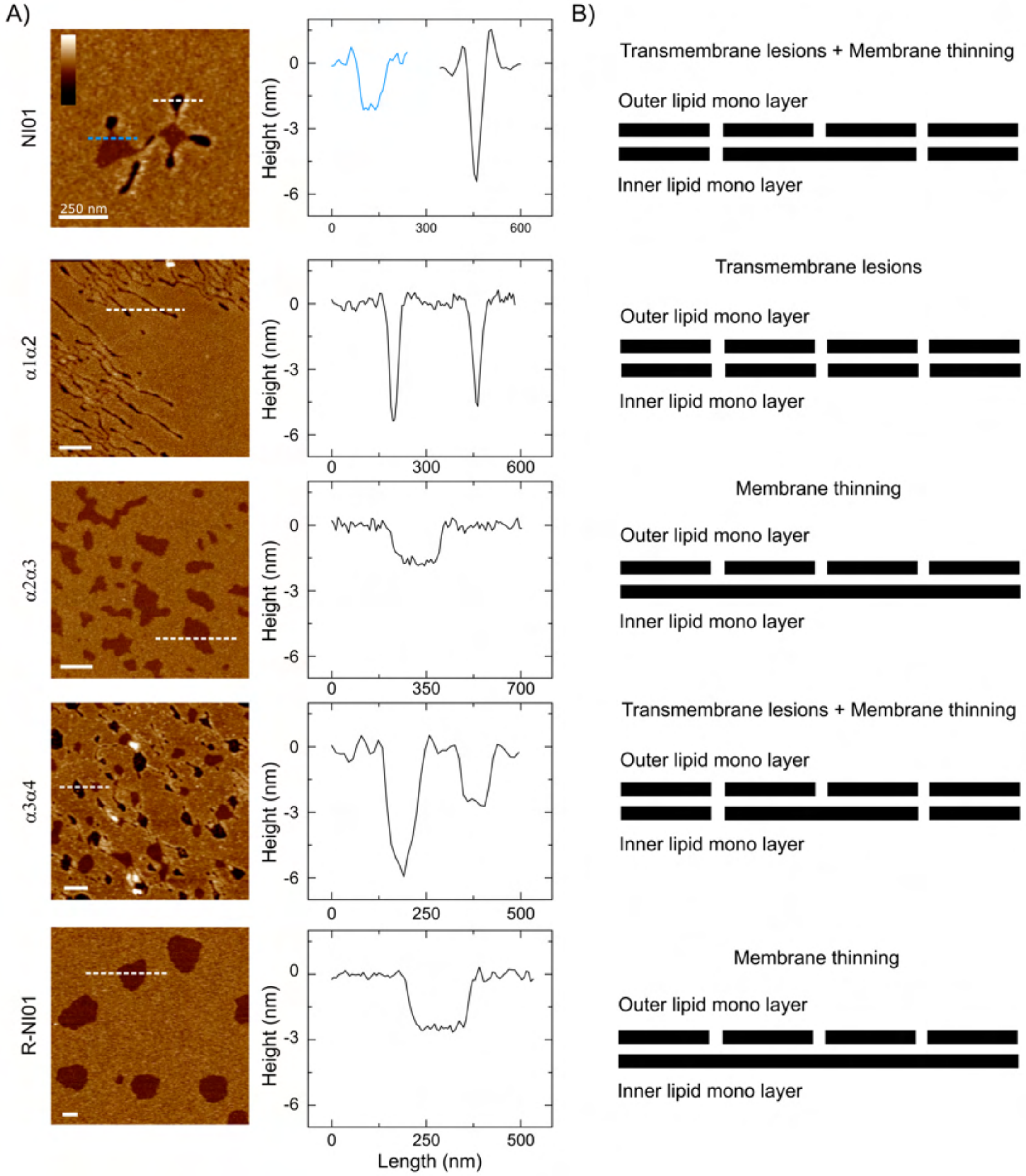


Figure S5: Outline for the suggested mechanisms of the bacteriocin-derived series of peptides. A) In liquid AFM images representing the topography of PC/PG (3:1) SLBs mimicking bacterial membranes treated with NI01, three substructures derived from the NI01 and the arginine mutant R-NI01. NI01 produces floral shaped topography that consists of bilayer spanning petal shaped transmembrane lesions (dark regions) and a membrane thinning patch in the center (light regions). B) Schematics representing the different mechanisms for each of the peptides as interpreted from the in-liquid AFM topography images.

### Device Operation

After assembly the chip is mounted and filled as previously described,[S1] briefly the fluids in the chip were controlled using 8 pressure ports and a single syringe pump module, 3 for the three OLA inlets (vesicle formation), 4 for the 8 perfusion inlets (each pressure port is split into two reservoir using external fittings) and 1 for the 3 valves. After mounting the device, the first step is filling the 3 dead-end valve control channels with milliQ water. Later, the main flow channel is filled with OA solution from the main outlet (outlet A) in the direction of outlet in the connector chip (outlet B). Shortly after the OLA reservoirs are connected and formation should start before the OA solution reaches to the OLA outlet and traps any remaining air. Once an air-free device is obtained, the experiment can commence. A valve is actuated by applying ∼200-500 mbar .

#### Vesicle Formation and Trapping

Vesicle formation and trapping is initiated by actuating valve1 and applying suction at the lower outlet A (10 µL/h). The GUV stream diverges from being exclusively directed into outlet B and towards the traps. Actuating V1 allows the suction stream to disregard the perfusion inlet stream and dedicates the volumetric flow onto the GUV stream in the connector chip and in turn increasing trapping efficiency without increasing the flow rate in the traps. The flow rate at the entrance of each chamber during trapping was calculated to be between 1 and 1.5 µL/h.

##### Peptide Perfusion

After vesicle entrapment and verifying traps are occupied with GUVs, valve1 is relaxed, valve2 is actuated and the suction tube is removed leaving outlet A open. This reduces the functioning platform circuitry to perfusion inlets and vesicle chambers, whilst excluding the OLA component and connector chip for the remainder of the experiment. The reservoir with the peptide mixture is exchanged with the connected IA reservoir for each chamber and the input pressure was adjusted to ∼10 mbar to drive the peptide solution into the chamber. During that process, the trapped GUVs are washed with the remaining IA buffer that predominate the tygon tubing and should amount to ∼32 µL. It was calculated that the application of ∼10 mbar at each perfusion inlet results in a flow rate at the entrance of each chamber of ∼10.7 µL/h. Acquisition is promptly initiated afterwards.

#### Data Analysis and Filtering

##### Analysis procedure

All experiments were analysed using a custom made python software package, the source code for which we have made available on github. The software receives 64 experimental videos each corresponding to a different field of view (FOVs) covering the full network of traps. These FOVs are grouped such that 8 consecutive FOVs constitute one experimental chamber. Each experimental video is analysed in turn, outputting a csv file containing intensity time traces for all GUVs identified. These csv files are collated to group GUV time traces by chamber. Subsequently, the GUV time traces within each chamber are ordered by the time at which a GUV Leakage Event occurs to produce heat plots as in Figure 2.

#### Description of analysis

First the start of the experiment is automatically identified. This is achieved by including a small concentration of tracer dye in the peptide solution perfused in the experiment. To identify the arrival of the drug, the software calculates the median intensity of the field of view over the course of the full experimental video. This approximates the background fluorescence due to the tracer dye. The time at which the background reaches half its final intensity is defined as the start of the experiment, t = 0. To aid detection of GUVs they are detected at the time just before a significant increase in the background intensity is observed. Detection is performed by first creating a binary image in which all pixels are classified as either foreground, 1 or background, 0 by Otsu's thresholding method.[S2] Here we exploit the fact that before the background dye has arrived, there are only two intensity classes in our image, one bright and one dark. The dye within the GUVs are the only source of bright pixels and all other pixels are background. Therefore, after thresholding the binary image delimits GUV pixels from background. Next, segmenting individual GUVs in the binary image is achieved using a 'euclidean distance transform' (EDT) on the binary image and then a peak detection function, both from the scikit image python library.[S3] The EDT calculates the euclidean distance of each foreground pixel from the nearest background pixel and assigns this value to the given foreground pixel to create a transformed image. For roughly circular foreground objects, peaks occur in the EDT at the object's centre (given it has the furthest distance to its nearest background pixel). Thus finding such peaks allows us to find the centre coordinates of individual GUVs across the FOV. About these centre coordinates, 30 x 30 pixel Regions of Interest (ROIs) are taken forward for analysis. For each subsequent frame in the experimental video, the analysis loops over all ROIs first finding pixels within GUVs by Otsu's method and thus the mean intensity over all pixels within a GUV. Eventually, after a LE, thresholding fails as the GUV intensity is no longer significantly greater than the background intensity. At this point the mean intensity across the whole

ROI is recorded for the remainder of the experiment. This procedure stops after either 600 mins or the end of the experimental video, whichever is first. The final background intensity is determined by finding the average intensity of the final 5 frames of the experimental video. This is subtracted from each GUV intensity profile. The profile is then normalised by its own maximum intensity.

#### Filtering

Various filtering procedures are required to distinguish spurious intensity changes from real Leakage Events. Post-hoc filtering procedures are run at the end of the analysis routine. One such spurious intensity drop occurs when a GUV escapes its trap and as such the intensity recorded in the corresponding ROI drops between consecutive frames. When this occurs, predominantly, the GUV reappears in the next frame in a position downstream of its initial ROI. These instances are detected by the software and the relevant intensity time traces removed from the sample. Where the GUV completely leaves the FOV, it is not possible to distinguish these events from a bursting LE and therefore it is not removed. As well as this, GUVs sometimes enter ROIs after the leakage event of the original occupier. These events are also removed from the final sample. A GUI was created to allow human oversight at several steps in the analysis procedure. For example, the GUI allows the user to identify that appropriate ROIs are found, view the single GUVs within their ROIs over the full experimental time and verify that thresholding occurs correctly.
